# Avian H7N9 influenza viruses are evolutionarily constrained by stochastic processes during replication and transmission in mammals

**DOI:** 10.1101/2022.04.12.488056

**Authors:** Katarina M. Braun, Luis A. Haddock, Chelsea M. Crooks, Gabrielle L. Barry, Joseph Lalli, Gabriele Neumann, Tokiko Watanabe, Masaki Imai, Seiya Yamayoshi, Mutsumi Ito, Yoshihiro Kawaoka, Thomas C. Friedrich

**Affiliations:** Department of Pathobiological Sciences, University of Wisconsin-Madison, Madison, WI, United States of America; Division of Virology, Institute of Medical Science, University of Tokyo, Japan; Department of Molecular Virology, Research Institute for Microbial Diseases, Osaka University; Center for Infectious Disease Education and Research (CiDER), Osaka University; The Research Center for Global Viral Diseases, National Center for Global Health and Medicine Research Institute, Tokyo, Japan

## Abstract

H7N9 avian influenza viruses (AIV) have caused over 1,500 documented human infections since emerging in 2013. Although wild type H7N9 AIV can transmit by respiratory droplets in ferrets, they have not yet caused widespread outbreaks in humans. Previous studies have revealed molecular determinants of H7N9 AIV virus host-switching, but little is known about potential evolutionary constraints on this process. Here we compare patterns of sequence evolution for H7N9 AIV and mammalian H1N1 viruses during replication and transmission in ferrets. We show that three main factors – purifying selection, stochasticity, and very narrow transmission bottlenecks – combine to severely constrain the ability of H7N9 AIV to effectively adapt to mammalian hosts in isolated, acute spillover events. We find rare evidence of natural selection favoring new or mammalian-adapting mutations within ferrets, but no evidence of natural selection acting during transmission. We conclude that human-adapted H7N9 viruses are unlikely to emerge during typical spillover infections. Our findings are instead consistent with a model in which the emergence of a human-transmissible virus would be a rare and unpredictable, though highly consequential, “jackpot” event. Strategies to limit the total number of spillover infections will limit opportunities for the virus to win this evolutionary lottery.

## Introduction

The potential emergence of a novel avian influenza virus in humans remains a significant public health threat ^1–3^. Despite recent advances in influenza surveillance and forecasting^4–6^, we still do not understand the evolutionary processes underlying the emergence of pandemic influenza viruses ^1,3^. H7N9 avian influenza viruses (AIV) naturally circulate in aquatic birds and have been endemic in poultry since the virus’s emergence in China in February 2013 ^7^. Since then, H7N9 viruses have spilled over into human populations, causing 1,568 confirmed infections with a case fatality rate approaching 40% across five epidemic waves ^8^. During the fifth and largest epidemic wave, some low-pathogenicity avian influenza (LPAI) H7N9 viruses acquired a novel motif in hemagglutinin (HA) that both facilitates systemic virus replication in chickens and enhances pathogenicity in mammals ^9–13^; these viruses are designated highly pathogenic avian influenza (HPAI) H7N9 viruses.

High pandemic potential is currently assigned to both H7N9 and H5Nx AIV ^14–21^. H7N9 viruses appear particularly threatening because, unlike H5N1 viruses, wild type H7N9 viruses can be transmitted between ferrets via respiratory droplet ^17,22,23^. In addition, H7N9 viruses are capable of binding human-type receptors, in which sialic acids are linked to galactose in an α(2,6) configuration ^17,24,25^. It is therefore unclear why there have been no documented cases of human-to-human transmission of H7N9 viruses ^26^. Several factors may contribute to poor H7N9 transmissibility in humans, including preferential binding to avian-type α(2,3) sialic acid receptors; reduced polymerase activity at 33°C, which approximates the human upper respiratory tract temperature; and suboptimal acid stability, impacting successful membrane fusion ^24,25,27–31^. Nonetheless, ongoing isolated human spillover infections remain concerning because they provide an opportunity for adaptation of H7N9 viruses to human hosts, laying the groundwork for future AIV outbreaks. To our knowledge, only one previous study examined H7N9 genetic diversity within hosts and reported lower levels of diversity in ferrets than in chickens ^24^. Natural selection can only act on the available genetic variation in a population, so limited H7N9 diversity in mammalian hosts could impede the efficiency of mammalian adaptation.

In 2017, Imai et al characterized the replication and pathogenicity of H7N9 AIV in ferrets ^17^. Using time series samples originally collected in that study, we performed whole-genome deep sequencing in technical duplicate and evaluated H7N9 evolutionary dynamics in seven ferret transmission events and in an additional nine infections not resulting in transmission. We compared the viral genetic diversity of these AIV in a mammalian system to the 2009 pandemic H1N1 virus in four ferret transmission events and one additional non-transmitting infection ^17,32^. While stochastic forces played a significant role in viral evolution, we found little evidence for H7N9 mammalian adaptation in ferrets. These observations suggest there is a high evolutionary barrier to the emergence of a H7N9 AIV capable of sustained spread in humans.

## Results

### H1N1 viruses transmit more frequently than H7N9 viruses in ferrets

We isolated and sequenced viral RNA (vRNA) from nasal washes collected from two previously published studies ^17,32^. Four of 5 donor ferrets infected with H1N1 virus (CA04) transmitted infection to a naive recipient ferret (80%). By comparison, 7 of 16 ferrets infected with H7N9 AIV transmitted to their recipients (43.7%) (**Figure 1**). These group sizes are small and, while the rate of transmission from H1N1-infected ferrets exceeded that of H7N9-infected ferrets, the difference was not significant (p=0.17; Mann-Whitney U).

**Figure 1:**
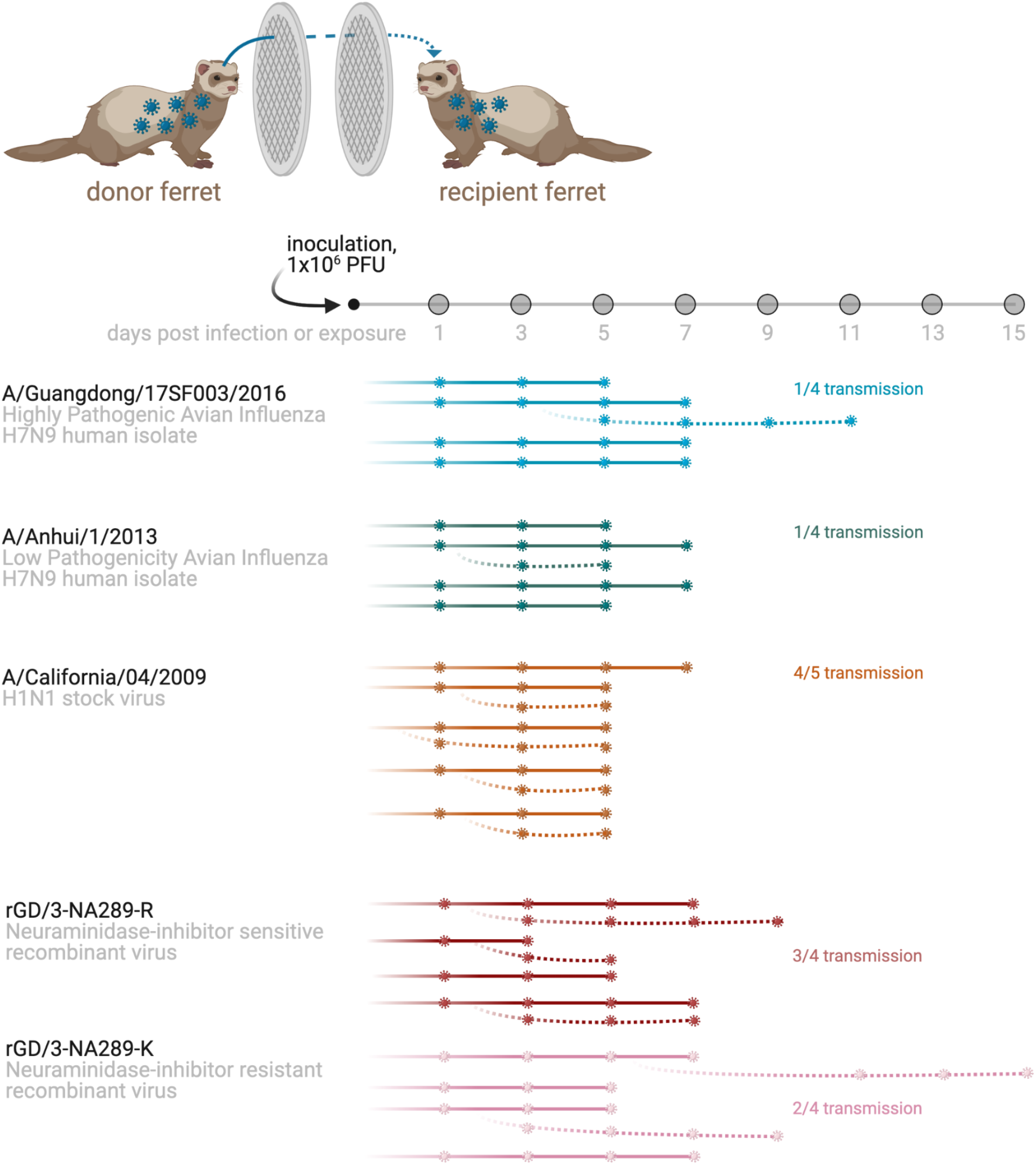
Overview of the experimental system and sampling timeline. Ferrets were inoculated intranasally with 10^6^ plaque-forming units (PFU) of a HPAI H7N9 isolate (A/Guangdong/17SF003/2016; blue), a LPAI H7N9 isolate (A/Anhui/1/2013; green), a H1N1pdm isolate (A/California/04/2009; orange), or recombinant GD/3 viruses (rGD3-NA289R; red, rGD3-NA289K; pink). One day after inoculation, one naive recipient ferret was paired with each donor ferret. Nasal washes were collected from donor (solid lines) and recipient (dotted lines) ferrets every other day up to 15 days post inoculation (DPI). Cartoon virions denote days on which live virus was detected by plaque assay and viral RNA was extracted for whole-genome sequencing. Nasal wash titers can be found in the GitHub repository accompanying this manuscript ^33^.

Rates of transmission varied across H7N9 virus subgroups (**Supplementary Table 1**). Transmission occurred in 1 of 4 ferret pairs whose donors were infected with either the LPAI isolate (A/Anhui/1/2013; “Anhui/1”) or the HPAI isolate (A/Guangdong/17SF003/2016; “GD/3”). The human GD/3 isolate contained neuraminidase-sensitive (NA-289R) and -resistant (NA-289K) subpopulations^17^, which were subsequently tested as recombinant GD/3 viruses in the 2017 Kawaoka study^17^. Two of four ferrets infected with the drug-resistant variant, rGD/3-NA289K, transmitted to recipient animals (50%), and three of four donors infected with the wild type recombinant variant, rGD3/NA289R, transmitted to the recipient (75%) (**Figure 1**).

### H7N9 within-host diversity is dominated by low-frequency iSNVs

We mapped sequencing reads to the inoculating virus consensus sequence and called within-host variants found in both technical replicates in ≥1% of reads (intersection iSNVs). iSNV frequencies from 1-99% were highly concordant between technical replicates (R^2^ = 0.993, **Supplementary Figure 1**). All coding region changes are reported using H7 numbering for the H7N9 viruses and H1 numbering for the H1N1 virus, consistent with the numbering schemes used in Nextstrain ^34^. We detected iSNVs in one or more viruses at 879 different genome sites, of which 490 were synonymous, 386 were nonsynonymous, and 3 were stop mutations. These stop mutations were found at low frequencies and were located at the terminal ends of coding regions (NP E292*, NS1 W203*, NEP Q119*).

Natural selection must act on existing genetic variation, so we first characterized the within-host diversity in the CA04 and H7N9 virus groups (Anhui/1, GD/3, rGD/3). The average number of iSNVs per ferret across all available time points varied significantly across virus groups (p=6.83×10^−10^; one-way ANOVA; **Figure 2a**). We found the fewest iSNVs per ferret in the CA04 group (n=9 ferrets), with an average of 24 iSNVs per ferret, ranging from 3-83. This is similar to the number of seasonal flu iSNVs reported in humans ^35^. The number of iSNVs in the rGD/3 group (n=13 ferrets, grouping both rGD/3 viruses) was also low, with an average of 13 per ferret, ranging from 1-43. We found more iSNVs in the ferrets infected with H7N9 biological isolates. Anhui/1 (n=5 ferrets) had an average of 152 iSNVs per animal, ranging from 85-195, while GD/3 (n=5 ferrets) had an average of 109 iSNVs per ferret, ranging from 27-142 across all timepoints. This level of within-host diversity is not unexpected in animals directly inoculated with a high-dose viral isolate ^36,37^. The number of iSNVs found in each animal fluctuates over time and often trends downward in GD/3 and Anhui/1 donor ferrets (**Supplementary Figure 2**).

**Figure 2:**
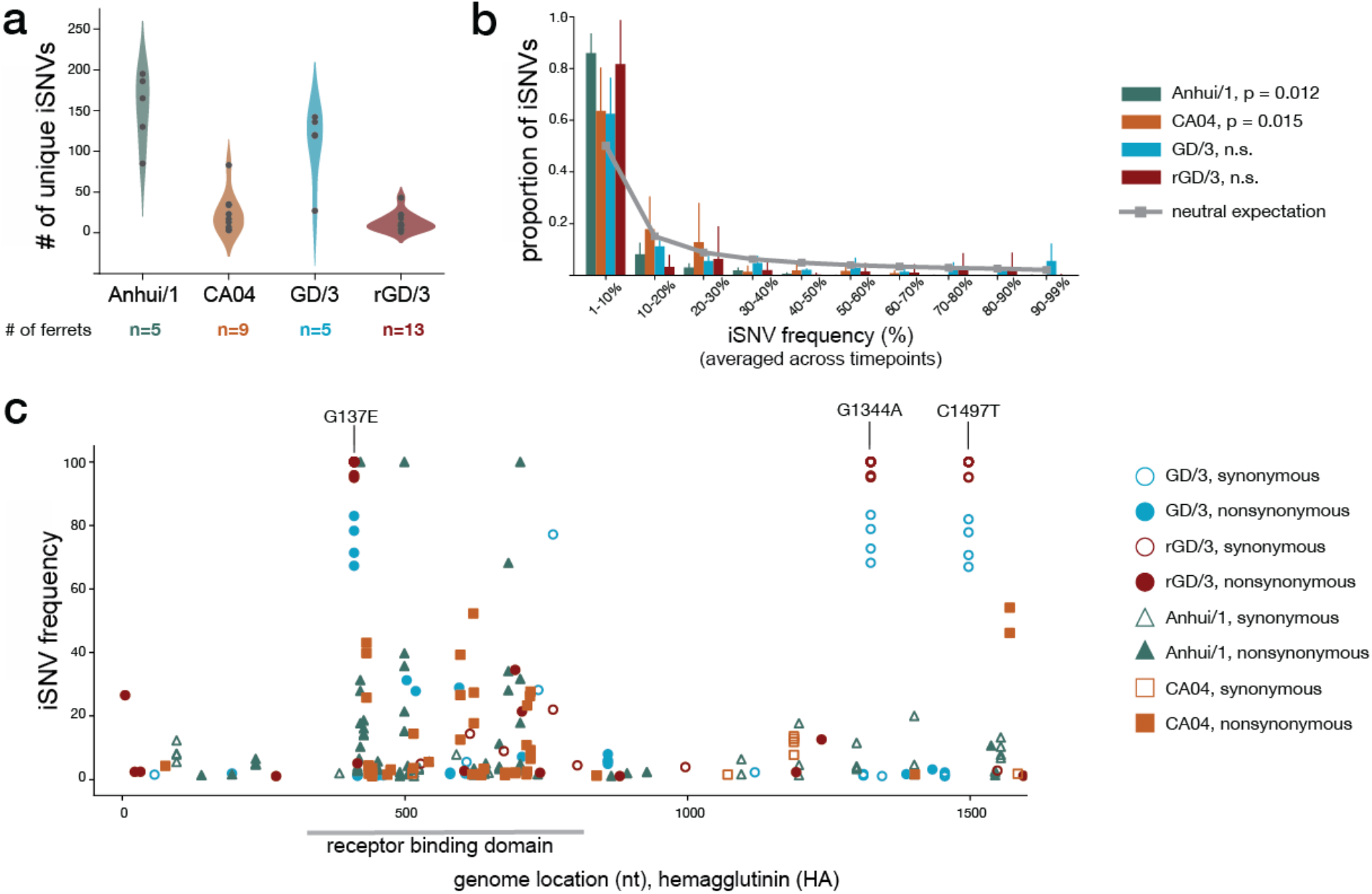
Frequency and location of intrahost single nucleotide variants. **a**. A violin plot showing the total number of iSNVs detected across all available time points plotted per virus group (p=6.83×10^−10^; one-way ANOVA). The individual data points denote the number of iSNVs per ferret. **b**. An iSNV frequency distribution showing the proportion of iSNVs detected per frequency bin. Error bars display variance (standard deviation) in the proportion of within-host iSNVs across ferrets The solid gray line denotes the expected proportion of variants in each frequency bin under a neutral model. Virus group distributions were compared to the neutral distribution using the Kolmogorov–Smirnov test. **c**. All iSNVs detected in hemagglutinin (HA) are plotted for GD/3 and rGD/3 iSNVs (circles), Anhui/1 (triangles), and CA04 (squares). Synonymous iSNVs are denoted with open symbols and nonsynonymous iSNVs with closed symbols. Three iSNVs found in multiple HPAI samples at high frequencies are labeled; G137E and two synonymous mutations at nucleotides 1,344 and 1,497. iSNVs in all other gene segments can be found in **Supplementary Figure 3**.

Most iSNVs were detected at <10% frequency (**Figure 2b**). Compared to expectations under a model of neutral evolution, low-frequency iSNVs were present in excess in Anhui/1, rGD/3, and to a lesser degree, GD/3 (see **Supplementary Table 2** for iSNV bin frequencies and variances within each group). This predominance of low-frequency iSNVs is consistent with viral population expansion and/or purifying selection acting within hosts. We used a Kolmonogorov-Smirnov test to compare the shape of the neutral distribution to the iSNV frequency distribution within each virus group. We found the Anhui/1 distribution (p=0.012) and the CA04 distribution (p=0.015) differed moderately from neutral and the GD/3 (p=0.787) and rGD/3 (p=0.052) distributions do not. The frequency, genome location, and impact on amino acid sequence (synonymous vs nonsynonymous) for iSNVs detected in HA across all ferrets is shown in **Figure 2c**. iSNVs in all other gene segments are plotted in **Supplementary Figure 3**.

### H7N9 viral populations are subject to purifying selection and genetic diversity is reduced following transmission

We used a common measure of nucleotide diversity, π, within individual ferrets to roughly assess signals of H7N9 viruses adapting to or diversifying within mammalian hosts. This summary statistic quantifies the average number of pairwise differences per nucleotide site among a set of viral sequences. In particular, we compared the nucleotide diversity at synonymous sites (πS) to nucleotide diversity at nonsynonymous sites (πN) to assess the evolutionary forces acting on each viral population. In general, πN/πS < 1 indicates that purifying selection is acting to remove deleterious mutations from the viral population, and πN/πS > 1 indicates that diversifying selection is favoring new mutations, which might be expected in the case of an avian influenza virus adapting to a mammalian host. When πN approximates πS, this suggests that allele frequencies are primarily determined by genetic drift, i.e., stochastic shifts in allele frequencies related to population size ^38^.

In most ferrets infected with H1N1 viruses, πS exceeded or was equal to πN (**Figure 3a**, orange). πS was significantly greater than πN in PB2, PA, and NA, and πN never significantly exceeded πS in any gene segment. These findings suggest that H1N1 virus populations in ferrets are shaped by mild purifying selection and genetic drift. This is expected for a mammalian virus replicating in a mammalian host, where most mutations away from a fit consensus are likely to be deleterious. Somewhat surprisingly, πN and πS comparisons gave similar results for H7N9 viruses. πS significantly exceeded πN in all genes apart from NA in the GD/3 group and all genes apart from NA and HA in Anhui/1 (**Figure 3a**, blue and green). Therefore, HPAI and LPAI H7N9 viruses are shaped by a combination of purifying selection and genetic drift, rather than diversifying selection as we might expect in the case of a virus adapting to a new host environment. The rGD/3 group had fewer iSNVs contributing to diversity measurements, but even still we found no evidence of diversifying selection as πN never significantly exceeded πS (**Figure 3a**, pink).

**Figure 3:**
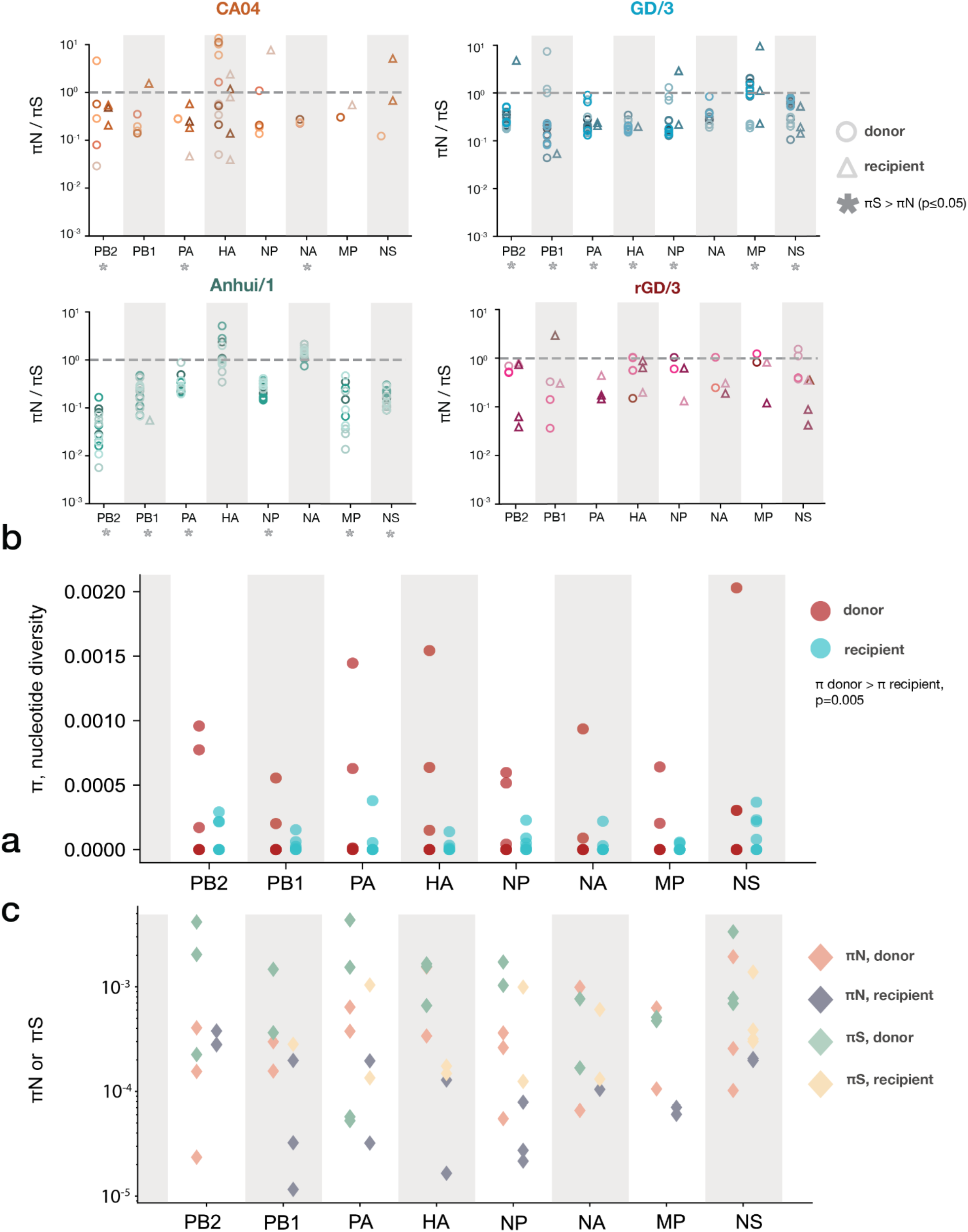
Patterns of viral genetic diversity within ferret hosts. **a**. πN / πS nucleotide diversity is plotted for each gene segment. Each data point represents a single ferret. Circles denote donor ferrets and triangles denote recipient ferrets. πN equal to πS (y=1) is represented with a dashed gray line. Gray stars denote instances when πS is significantly greater than πN. **b**. Genewise nucleotide diversity is plotted for all H7N9 transmission pairs (Anhui/1, GD/3, and rGD/3). The donor ferrets are shown in brick red and the recipient ferrets are shown in aqua blue. Nucleotide diversity did not significantly differ between donor and recipient ferrets in any single gene segment, but is significantly lower following transmission in recipient ferrets when assessing the entire genome (p=0.005; paired t-test). **c**. πN and πS in the H7N9 donors and recipients are plotted for each gene segment. πN and πS in the donor ferrets are denoted by the salmon and light green diamonds, respectively. πN and πS in the recipient ferrets are denoted by the dark blue and yellow diamonds, respectively. Similar data are plotted for H1N1 transmission pairs in **Supplementary Figure 4**.

We also compared nucleotide diversity in donor-recipient pairs before and after transmission. We found genome-wide nucleotide diversity (π) did not significantly differ between donor and recipient ferrets in the H1N1 group (**Supplementary Figure 4a**, p=0.125, paired t-test). However, in the H7N9 group, π in the donor ferrets was significantly greater than recipient ferrets (**Figure 3b**, p=0.005; paired t-test). It is clear that, overall, H7N9 genetic diversity is lost during transmission. As we have done previously ^36,39^, we looked for selective sweeps by comparing the change in πN and πS in each gene segment for paired donor and recipient ferrets. πN and πS at the gene level did not differ significantly between donor and recipient ferrets. This was true across all H1N1 transmission pairs (**Supplementary Figure 4b**) and all H7N9 transmission pairs (**Figure 3c**). These findings suggest that during transmission of these viruses, genetic diversity was purged equally across gene segments with no evidence for a selective reduction in diversity of any particular segment.

### Airborne transmission results in a dramatic shift of iSNV frequencies

We took advantage of time series data to follow individual iSNVs within donors and following transmission into recipient ferrets. Strikingly, the frequency of an H7N9 iSNV in a donor predicts neither its likelihood of transmission nor its frequency post-transmission. For example, one iSNV that encodes a glycine-to-glutamic-acid substitution at HA position 137 (HA G137E) in the GD/3 transmission pair was present at 81% at 1 DPI in the donor ferret and decreased to a sub-consensus frequency (39.3%) by 7 DPI. Despite this marked downward trend in the donor animal, G137E was transmitted to the recipient at 5 DPI and remained at ≥99% from this time point onward (**Figure 4**). Conversely, a mutation in the matrix gene encoding an arginine-to-lysine substitution at position 210 in M1 (R210K) was never detected in the donor ferret above 1%, yet was nearly fixed (97.5%) at the first time point post-infection in the recipient. Interestingly, M1 R210K then decreased in frequency in the recipient and was found at 54.5% at 9 DPI. We observed similar patterns in synonymous variants. For example, a synonymous A-to-G change at nucleotide (nt) 2,037 in the polymerase basic protein 1 (PB1) gene was never found above 6% frequency in the donor ferret, but was nearly fixed immediately following transmission, and again decreased in frequency to 57.57% by 9 DPI in the recipient ferret. It is important to note that down trending iSNV frequencies may be influenced by stochastic fluctuations or genetic drift within the population and are not necessarily a function of fitness.

Even in the case of amino acid substitutions that may be adaptive in humans, variant frequencies were not maintained across the transmission event. For instance, a valine-to-isoleucine substitution at position 219 in M1, which may play a role in avian influenza virus adaptation to mammals ^40^, increased in frequency from 34.7% to 84.3% in one donor, but nonetheless failed to transmit to the recipient. M1 V219I then appeared to arise *de novo* in the recipient ferret, suggesting that positive selection may act on individual sites within individual hosts, despite a lack of evidence for positive selection at the gene level at the time of transmission. A similar trend was observed for an aspartic-acid-to-asparagine substitution at position 701 in PB2, a mutation associated with enhanced replication in mammals ^41–43^. No variants consistently increased in frequency over time in the Anhui/1 or rGD/3 groups. Unlike iSNV dynamics in the H7N9 transmission events, which resulted almost exclusively in fixation or loss of iSNVs in the recipient ferrets, eight iSNV sites in the H1N1 CA04 donor ferrets remained polymorphic at intermediate frequencies immediately following transmission (e.g. HA D127E and S183P) (**Figure 4c**).

**Figure 4:**
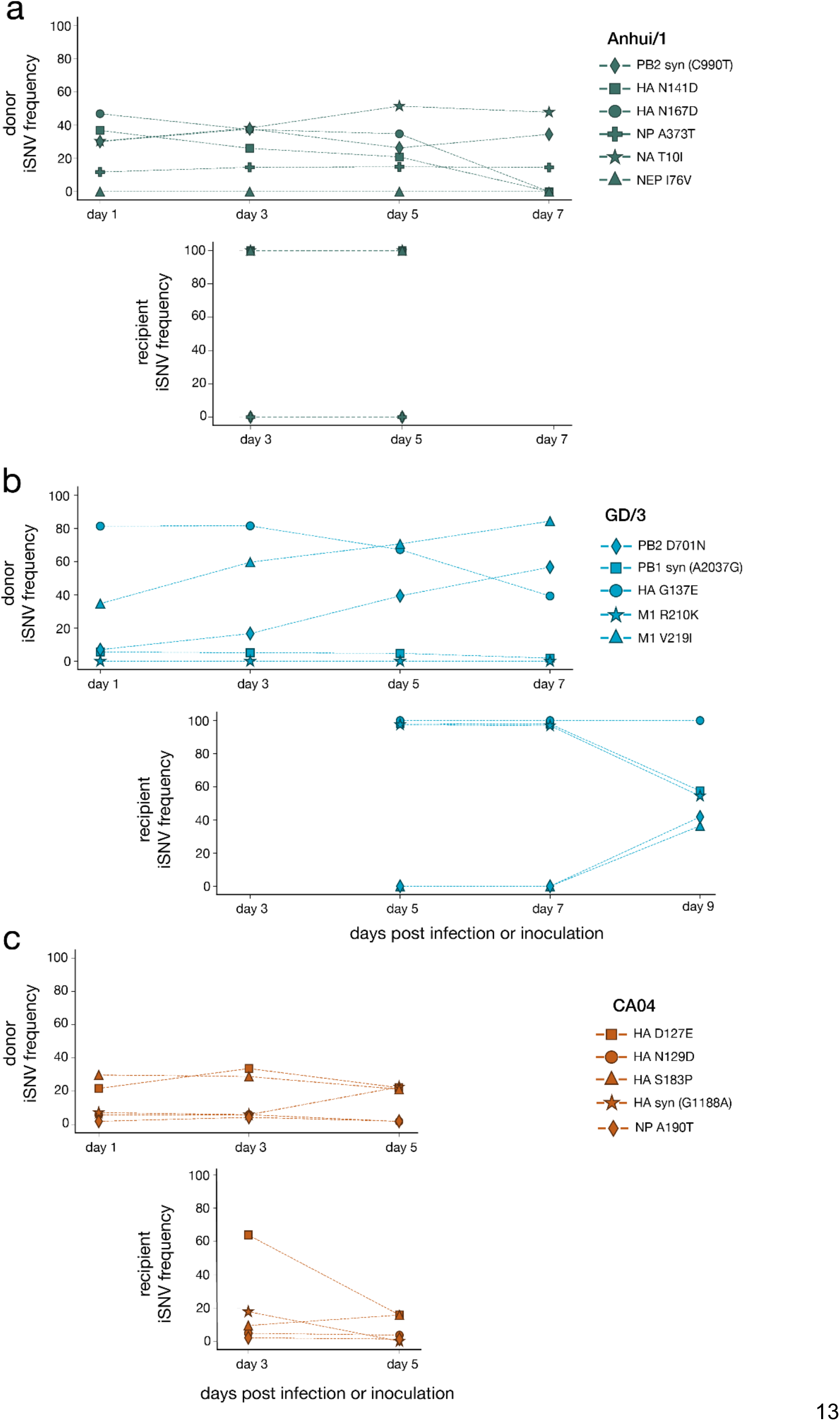
Frequency dynamics of iSNVs across the transmission event. The frequencies of representative iSNVs are plotted over time in donor ferrets (top plot) and following transmission into the associated recipient ferret (bottom plot) in the **a**. Anhui/1 transmission pair, **b**. GD/3 transmission pair, and **c**. CA04 transmission pairs. Each iSNV is plotted as y=0 at time points when it is not detected ≥1% frequency and is absent at timepoints when no viral RNA was recovered for deep sequencing. We did not plot iSNV frequencies beyond day 9 in the recipient ferret.

These results highlight how airborne transmission can dramatically alter the frequency of influenza virus variants across hosts. While the vast majority of variants are lost at the time of transmission, we observed that deleterious mutations can be transmitted and putatively adaptive ones may not. These observations suggest that natural selection at the time of H7N9 transmission is negligible. Additional work to characterize the fitness benefit or cost of individual mutations are needed to determine the full range of evolutionary forces acting within individual hosts.

### Airborne transmission of H7N9 viruses in ferrets is characterized by a very narrow transmission bottleneck

Narrow transmission bottlenecks, in which a very small number of viruses found a new infection, cause a founder effect and purge most low-frequency iSNVs, regardless of their fitness ^44,45^. Conversely, wide transmission bottlenecks allow more viruses to initiate infection, reducing the chance that beneficial or rare variants are lost. The vast majority of iSNVs detected in all H7N9 and H1N1 donor ferrets were lost during transmission and were not found in recipients. However, a very small number of iSNVs in the Anhui/1 and GD/3 donor ferrets were transmitted and fixed (>99% frequency) in the recipient ferret (**Figure 5a**).

**Figure 5:**
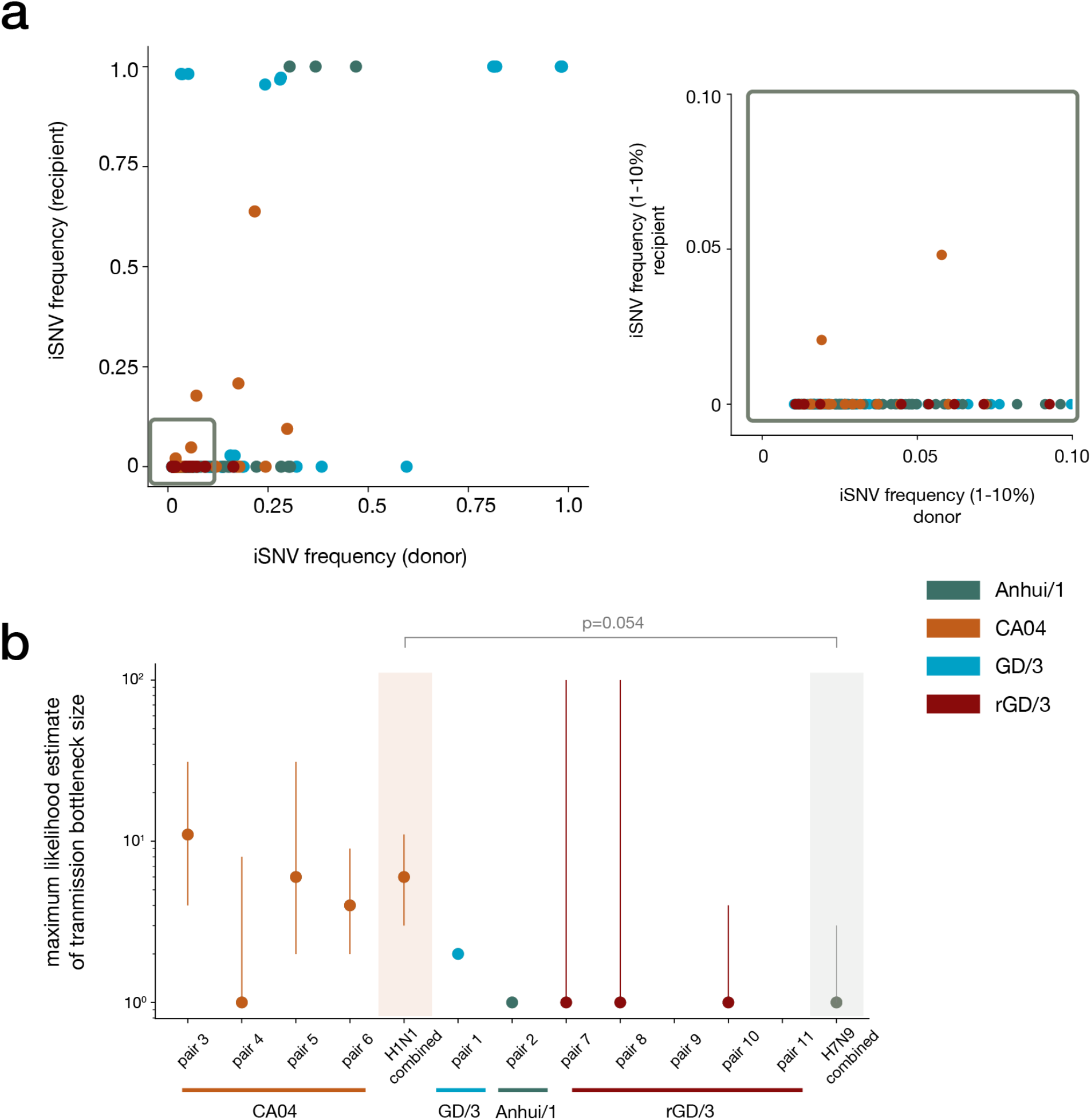
H1N1 and H7N9 transmission bottlenecks in ferret donor-recipient pairs. **a. “**TV plots” showing intersection iSNV frequencies in all 11 donor-recipient pairs. The inset box highlights low-frequency iSNVs (1-10%). Colors denote virus groups. **b**. Maximum likelihood estimates for mean transmission bottleneck size with 95% confidence interval in individual donor-recipient pairs. Bottleneck sizes could not be estimated for two pairs (rGD/3 pair 9 and pair 11) because there were no polymorphic sites detected in the donor. The combined H1N1 estimate was calculated using pairs 3, 4, 5 and 6. The combined H7N9 estimate was calculated using pairs 1, 2, 7, 8 and 10 (p=0.054; unpaired t-test).

To infer transmission bottleneck sizes, we applied the beta-binomial inference method ^46^ to estimate the number of transmitted viruses that could account for the pattern of iSNVs observed immediately before and after transmission for each pair. These estimates suggest that fewer than 11 viruses initiated infection in all recipient ferrets. The combined maximum likelihood estimate for the mean transmission bottleneck size for the CA04 (H1N1) pairs was 6 (n=4 pairs; 95% CI: 3-11; **Figure 5b**). We evaluated seven transmission events in the H7N9 group: one Anhui/1 pair, one GD/3 pair, and five rGD/3 pairs. However, two of the rGD/3 transmission events (pairs 9 and 11) were uninformative as the donor had no detectable polymorphic sites. The combined maximum likelihood for the mean transmission bottleneck size among the remaining H7N9 pairs was 1 (95% CI: 1-3; **Figure 5b**). The combined H1N1 transmission bottleneck estimate is larger (looser) than the combined H7N9 estimate, although with only nine transmission pairs informing these estimates, this difference did not reach statistical significance (p=0.054; unpaired t-test).

### H7N9 iSNVs arising in ferrets are represented equally in global H7N9 viruses collected from both birds and humans

Each H7N9 infection in a human represents a unique avian spillover event. If selection is strong at a given site in the genome, then we might observe mutations at that site in multiple independent infections. To look for such a signal, we compared nonsynonymous iSNVs detected in this study (n=262) to nonsynonymous SNPs found at the most distal nodes in Nextstrain’s H7N9 phylogenetic tree (n = 2,071) ^34^. This tree was created from publicly available H7N9 sequences collected from birds (n=621) and humans (n=1,023). Among the list of variants shared between our data and Nextstrain’s, we looked for those that were proportionally overrepresented in sequences from humans. We excluded iSNVs detected in the rGD/3 samples because the inoculum was near clonal and few iSNVs were detected in ferrets.

Considering all iSNVs we detected, around half (46.6%, 77/165) of the Anhui/1 iSNVs and 36.1% (35/97) of the GD/3 iSNVs can be found in at least one bird or human H7N9 sequence on Nextstrain. Of the 77 Anhui/1 iSNVs, 49 were in human sequences, 9 were in bird sequences, and 19 were found in both. Of the 35 GD/3 iSNVs, 20 were in human sequences, 8 were in bird sequences, and 7 were found in both. A complete summary of iSNVs and their respective occurrences in human-or bird-derived sequences from all gene segments can be found in **Supplementary Table 3**. As a group, neither the Anhui/1 variants nor the GD/3 variants were significantly enriched in human sequences compared to bird sequences (GD/3 p = 0.052, Anhui p = 0.049; Fisher’s exact test).

We plotted the number of occurrences of each of our iSNVs in bird and human sequences in **Figure 6**. Four Anhui/1 iSNVs were significantly enriched in bird sequences (HA L235Q, HA D289N, NA V22I and NS R44K). Two iSNVs were enriched in mammalian sequences; PB2 K627E in GD/3 (discussed further below) and PB2 D701N in Anhui/1, which is linked to mammalian adaptation. We also identified a few putative mammalian-adapting mutations that arose sporadically in ferrets, but were not enriched in human surveillance sequences. These mutations included PB2 K562R ^47^, HA A143T ^48^, MP V219I ^40^.

**Figure 6:**
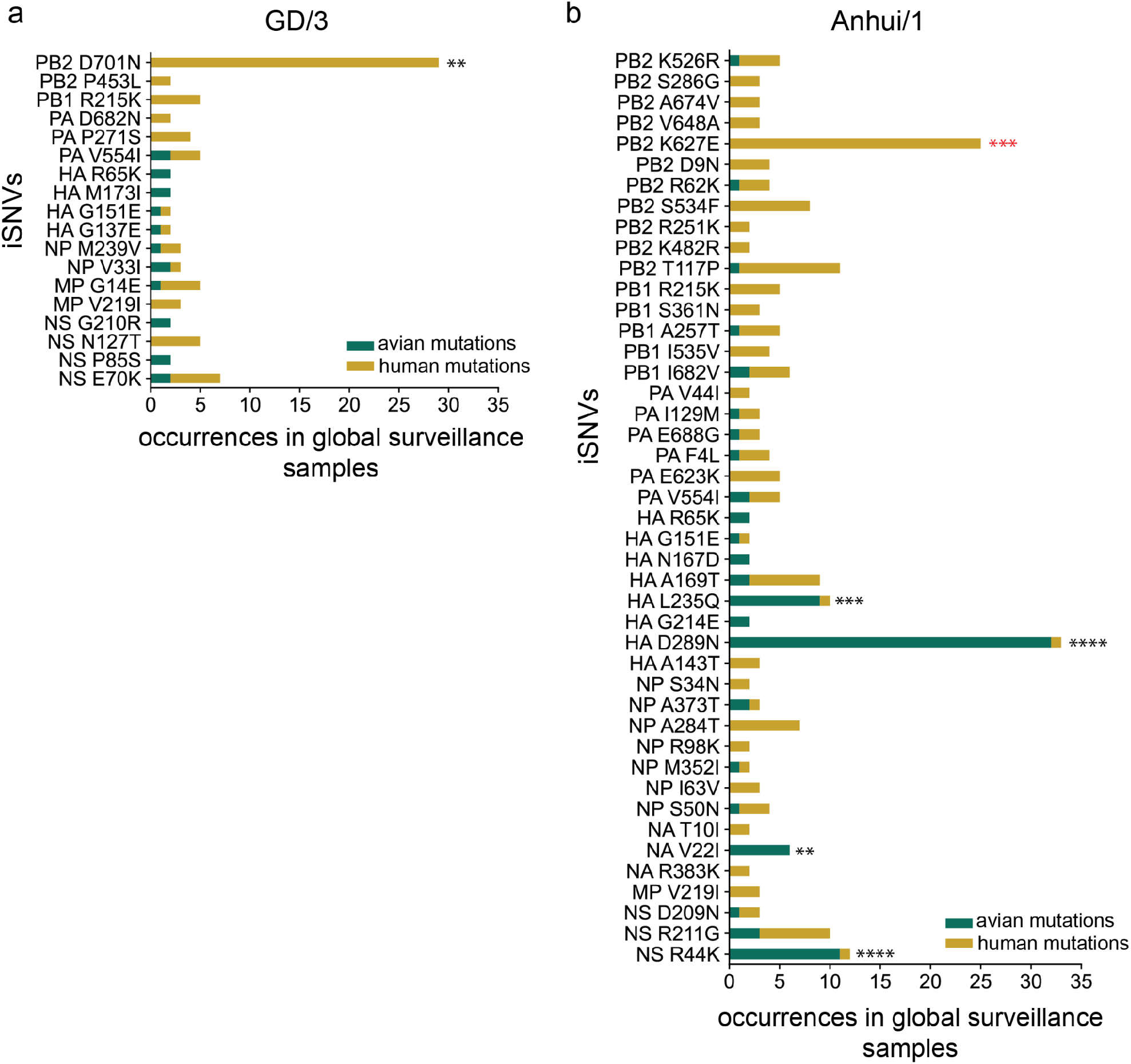
iSNVs found in H7N9 global surveillance sequences. Occurrences of iSNVs in H7N9 global surveillance sequences in the (**a**) GD/3, (**b**) and Anhui/1 datasets. This figure displays iSNVs with more than 2 occurrences. A plot with all iSNVs can be found in **Supplementary Figure 5**. Fisher’s exact test was used to assess for significant enrichment in human or bird sequences and these results are shown with asterisks (* p = 0.05 - 0.01, ** p = 0.01 - 0.001, *** p = 0.0001 - 0.0000).

A glutamic-acid-to-lysine change at residue 627 in PB2 (E627K) is a key mutation previously shown to improve polymerase processivity in mammalian hosts ^41–43^. The Anhui/1 isolate’s consensus sequence, which we used as our reference, already contained a lysine at this residue. Therefore, we report this iSNV as a lysine-to-glutamic-acid change (K627E) above. Importantly, although this iSNV met criteria for inclusion in this surveillance analysis, we only detect it (the glutamic acid change) in a single ferret, at a single point, near 1% frequency (**Figure 6b**, red asterisks).

## Discussion

The evolutionary pathways by which avian influenza viruses might adapt to cause widespread outbreaks in humans are poorly defined. Our study examined the viral dynamics of wildtype LPAI and HPAI H7N9 viruses in a ferret model, a well-studied mammalian system which closely recapitulates human respiratory physiology and transmission ^49^. We found that H7N9 viruses in ferrets are subject to mild purifying selection and that as a group, HPAI H7N9 iSNVs are equally represented in bird and human surveillance sequences. These findings are consistent with a virus that is already sufficiently fit for replication in humans and is not under strong selective pressure in mammalian systems. However, our results shed light on several significant barriers to the generation, selection, and, in particular, onward transmission of such mutations. We speculate that short infection times, purifying selection, and narrow, non-selective transmission bottlenecks combine to limit the capacity of H7N9 viruses to adapt during typical spillover infections.

Some have theorized that the rate-limiting step in viral host switching is not the generation of adaptive variants within hosts, but the successful transmission of these variants between them ^16^. Our data support this hypothesis. We detected multiple iSNVs in donor ferrets putatively linked to mammalian adaptation (PB2 K526R, PB2 D701N, HA A143T, and M1 V219I) or enriched in human surveillance sequences (PB2 D701N) that were not transmitted onward. Indeed, the vast majority of iSNVs arising in ferrets were lost during transmission. This is the result of an extremely narrow transmission bottleneck—our estimates indicate that new infections were founded by 1-3 H7N9 viruses. Our quantitative results thus confirm a speculation previously made by Zaraket et al. in a study of LPAI H7N9 transmission in ferrets ^24^.

If selection does not act efficiently during transmission, any mutations on a transmitted genome are likely to become dominant in the recipient viral population. These variants, however, may not be the fittest variants present in the donor population, so transmission may reduce overall viral fitness. We saw two examples of this when M1 V219I and PB2 D701N, both previously linked to mammalian adaptation ^40^, were lost at the time of transmission, and arose *de novo* once again in the recipient ferret. Therefore, when transmission bottlenecks are narrow and selection at the time of transmission is negligible, common, mildly deleterious variants may become fixed and low-frequency adaptive variants are very likely to be lost, ultimately slowing the pace of viral adaptation. These evolutionary barriers imposed by narrow transmission bottlenecks may not be unique to H7N9 viruses. Our recent study evaluating the adaptive potential of H5N1 viruses suggested that H5N1 evolution is also limited by the loss of genetic variation resulting from airborne transmission ^50^. However, that study involved inferences based on unlinked human and avian infections. Here the ferret model system allows us to unambiguously analyze linked donors and recipients and quantify bottleneck sizes.

Despite the success of mass poultry immunizations with the H5/H7 bivalent vaccine, H7N9 viruses are likely to continue to evolve and sporadically spillover into humans. H7N9 remains common in poultry and large populations of unvaccinated duck and wild waterfowl may serve as reservoirs for ongoing adaptation and reassortment of HPAI H7N9 viruses ^51^. Furthermore, Wu et al characterized H7N9 viruses collected at live poultry markets and farms between 2017 and 2019 and found evidence for accelerated evolution away from the vaccine strain in the 2018-2019 swabs ^52^. Our study suggests that H7N9 is unlikely to acquire enhanced human transmissibility during a single infection. However, this risk is additive and may become non-negligible with an increasing number of human spillover infections. This emphasizes the importance of population health interventions to reduce opportunities for avian viruses to spill over into humans and, even more so, the opportunity for avian and mammalian viruses to co-infect a single host. These interventions are reviewed in full in the China-WHO Joint Mission on Human Infection with Avian Influenza A(H7N9) Virus ^53^ and include, but are not limited to, continued poultry vaccination, culling, poultry movement restrictions, distancing at live animal markets, and others ^54^.

We speculate that the evolutionary processes and the patterns of selection acting on wildtype avian viruses in a mammalian system are distinct from those acting on reassortant or engineered avian viruses. Ferguson et al ^55^ and Sobel-Leonard et al ^56^ have previously described the concept of viral fitness landscapes, which are defined by “fitness peaks” and “fitness valleys’’ resulting from unique combinations of virus and host genotype interactions. The topography of this landscape is expected to change with shifting host immune environments, epistatic interactions, new reassortant genotypes, etc. The fitness peaks, areas of high viral fitness, on this landscape are occupied by viruses like seasonal H1N1 in a human (a wild type mammalian virus in a mammal) and H7N9 in a chicken (a wild type avian virus in a bird). These viruses are already well adapted to their hosts and have limited nearby evolutionary space to become fitter; that is, mutations in these viruses will tend to be deleterious, moving the virus away from a local fitness peak. Such viral populations are likely to be characterized by purifying selection and genetic drift because any new mutation is unlikely to possess a large enough selection coefficient to be positively selected in the setting of an acute infection. Perhaps counter-intuitively, our results indicate that H7N9 avian viruses are already relatively fit in ferrets, a mammalian host. We saw no evidence of adaptive evolution within hosts, and a regime in which genetic drift and purifying selection dominate. Altogether this suggests that most mutations in H7N9 viruses replicating in ferrets are deleterious, as we would expect for viruses near local fitness maxima.

Interestingly, we also did not find evidence for selection during transmission of avian H7N9 viruses in mammals. This contrasts with our previous studies in which ferret transmission of genetically engineered H5N1 and “1918-like” H1N1 avian influenza viruses was associated with selective sweeps acting on HA ^39,57^ In these sweeps, selection appeared to favor transmission and/or replication of only a subset of HA sequences in recipient ferrets, as evidenced by sharp decreases in genetic diversity in HA, but not other gene segments. We posit these engineered viruses resemble hypothetical “transitional states” distant from local fitness maxima. For such viruses, many new mutations may confer fitness advantages and be positively selected within hosts and/or swept to fixation during transmission. We might expect such unfit viruses to be unstable and therefore likely transient in nature. However, selective sweeps between hosts or rapid diversification within a host may be “evolutionary signatures” that indicate viruses with heightened pandemic potential. Importantly, surveillance approaches aimed at detecting evolutionary signatures of within-and/or between-host selection would be agnostic to AIV subtype, genetic background, and are less likely to be confounded by epistasis than traditional approaches that query lists of mutations of concern. Such sequence-agnostic approaches could therefore provide an important complement to traditional risk assessments for avian influenza viruses, particularly for subtypes for which there is little data on the phenotypic impact of specific mutations.

Like most ferret studies, the results of these experiments are limited by relatively small sample sizes and the biological differences between ferrets and humans. Ferrets are the most relevant animal model system for studying respiratory infections; however, there are anatomical, physiological, and immunological differences between ferrets and humans, highlighted by the fact that H7N9 AIVs are transmitted between ferrets but are not known to do so between humans ^58^. Accordingly, there may be fewer or different evolutionary pressures acting on the H7N9 viruses in ferrets compared to humans. We also included clonal, recombinant viruses (rGD/3), which, as stated previously, have less diversity than viral isolates and will thus be subject to different evolutionary forces. In addition, direct inoculation of donor ferrets does not fully recapitulate a human spillover infection. In particular, high-dose inoculation with a biological isolate may allow a greater number of more diverse genomes to establish infection than in natural infections. Patterns of genetic diversity might differ in the case of H7N9 transmitting directly from a bird to a human. Our results should be corroborated by further investigations, including natural spillover infections if possible, as well as targeted virological and epidemiological research ^59^.

Assessing zoonotic risk and adaptive potential of AIV remains a critical public health challenge. By characterizing patterns of within-host diversity, quantifying the stringency and patterns of selection acting on typical transmission bottlenecks, identifying the fate of known adaptive mutations within individuals and across transmission events, and characterizing typical and non-typical evolutionary signatures, we can continue to assemble an understanding of AIV evolution. We hope these methods may be applicable to other zoonotic respiratory viruses, including SARS-CoV-2, in order to better assess their ongoing adaptive potential.

## Materials and methods

### Ferrets transmission experiments & sample collection and availability

No new transmission experiments were performed as part of this study. We took advantage of nasal wash samples collected from ferrets participating in a 2017 study conducted by Imai and colleagues to assess the transmissibility of H7N9 viruses in ferrets ^17^. In this previously-described study, four groups of four ferrets were directly inoculated with various H7N9 viruses (1 × 10^6^ PFU) and one group of two ferrets was infected with an H1N1pdm virus for comparison (inoculated or donor ferrets). Samples from three of the four total H1N1 pairs were derived from a separate and similar study by the Kawaoka group ^32^. The H7N9 viruses used in this study included a high-pathogenic human isolate – A/Guangdong/17SF003/2016 (“GD/3”), two recombinant viruses which have an arginine or lysine at position 289 (H7 numbering) to confer neuraminidase-inhibitor sensitivity or resistance, respectively, on the background of the GD/3 consensus sequence – rGD/3-NA289R and rGD/3-NA289K (“rGD/3”), and a low-pathogenic H7N9 virus – A/Anhui/1/2013 (“Anhui/1”). The H1N1 comparator group was inoculated with a representative 2009 pandemic virus – A/California/04/2009 (“CA04”).

Four (GD/3, rGD/3-NA289R, rGD/3-NA289K, Anhui/1) or six (CA04) serologically-confirmed naive ferrets (recipient ferrets) were placed in enclosures adjacent to the donor ferret (separated by ~5cm) on day 2 post inoculation. Pairs of ferrets were individually co-housed in adjacent wireframe enclosure which allow for spread of virus by respiratory droplet, but not by direct or indirect (via fomite) contact. Nasal washes were collected from donor ferrets on day 1 after inoculation and from recipient ferrets on day 1 after co-housing, and then every other day (for up to 15 days) for virus titration. Virus titers in nasal washes were determined by plaque assay on MDCK cells. Viral RNA was available for isolation from nasal wash samples collected from donor ferrets on days 1, 3, 5 and 7 post-infection and from recipient ferrets on days 3, 5, 7, 9, 11, 13, and 15 post-infection.

### Viruses

A/Guangdong/17SF003/2016 was propagated in embryonated chicken eggs to prepare a virus stock after being isolated from a fatal human case treated with oseltamivir ^60^. We sequenced this inoculum and plot iSNVs in **Supplementary Figure 6**. A/Anhui/1/2013 was also propagated in embryonated chicken eggs after being isolated from an early human infection ^22^. A/California/04/2009 was propagated in MDCK cells and was originally obtained from the Centers for Disease Control (CDC) ^61^. Recombinant viruses, rGD3-NA289K and rGD3-NA289R, were generated by plasmid-based reverse genetics as previously described ^62^.

### Template preparation

Total nucleic acids including viral RNA (vRNA) were extracted from nasal washes and were reverse transcribed using SSIV VILO (Invitrogen, USA) and the Uni12 primer (AGCAAAAGCAGG) in a total reaction volume of 20 µl. The complete reverse transcription protocol can be found here: https://github.com/tcflab/protocols/blob/master/VILO_Reverse_Transcription_h7n9_GLB_2019-02-15.md.

Single-stranded cDNA was used as a template for PCR amplification to amplify all eight genes using segment specific primers using high-fidelity Phusion 2X DNA polymerase (New England BioLabs, Inc., USA). Primer sequences are available in the GitHub repository accompanying this manuscript ^33^. PCR was performed by incubating the reaction mixtures at 98°C for 30 s, followed by 35 cycles of 98°C for 10 s, 51 - 72°C depending on gene segment for 30 s, 72°C for 120 s, followed by a final extension step at 72°C for 5 min. The complete PCR protocol, including segment-specific annealing temperatures and primer sequences, can be found here: https://github.com/tcflab/protocols/blob/master/Phusion_PCR_h7n9_GLB_2019-02-21.md. PCR products were separated by electrophoresis on a 1% agarose gel (Qiagen, USA). The bands corresponding to full-length gene segments were excised and the DNA was recovered using QIAquick gel extraction kit (Qiagen, USA). To control for RT-PCR and sequencing errors, especially in low-titer samples, all samples were prepared in complete technical replicate starting from vRNA ^63,64^. We sequenced samples with low or no coverage, typically from low-titer samples, a third time and merged sequencing reads with the first two replicates to minimize coverage gaps.

### Deep sequencing

Gel-purified PCR products were quantified using Qubit dsDNA high-sensitivity kit (Invitrogen, USA) and were diluted in an elution buffer to a concentration of 1 ng/µl. All segments originating from the same samples with a non-zero concentration as determined by hsDNA Qubit (Invitrogen, USA) were pooled equimolarly and these genome pools were again quantified by Qubit. Each equimolar genome pool was diluted to a final concentration of 0.2 ng/µl (1 ng in 5 µl volume). Each sample was made compatible for deep sequencing using the Nextera XT DNA sample preparation kit (Illumina, USA). Specifically, each sample or genome was enzymatically fragmented and tagged with short oligonucleotide adapters, followed by 15 cycles of PCR for template indexing. Individual segments with undetectable concentrations by Qubit dsDNA were tagmented and indexed separately to maximize recovery of complete genomes.

Samples were purified using two consecutive AMPure bead cleanups (0.5x and 0.7x) and were quantified once more using Qubit dsDNA high-sensitivity kit (Invitrogen, USA). If quantifiable at this stage, independent gene segments were pooled into their corresponding genome pools. The average sample fragment length and purity was determined using Agilent High Sensitivity DNA kit and the Agilent 2100 Bioanalyzer (Agilent, Santa Clara, CA). After passing quality control measures for loading the sequencing machine, genomes were pooled into six groups of ~30 samples, which were sequenced on independent sequencing runs. Libraries of 30 genomes were pooled in equimolar ratios to a final concentration of 4 nM, and 5 µl of each 4 nM pool was denatured in 5 µl of freshly diluted 0.2 N NaOH for 5 min. Denatured pooled libraries were diluted to a final concentration of 16 pM, apart from the first library which was diluted to 12pM, with a PhiX-derived control library accounting for 1% of total DNA loaded onto the flow cell. A total of 600 µl of diluted, denatured library was loaded onto a 600-cycle v3 reagent cartridge (Illumina, USA). Average quality metrics were recorded, reads were demultiplexed, and FASTQ files were generated on Illumina’s BaseSpace platform ^65^.

### Sequence data analysis – quality filtering and variant calling

FASTQ files were processed using custom bioinformatic pipelines, available on GitHub https://github.com/tcflab/Sniffles2. Briefly, read ends were trimmed to achieve an average read quality score of Q30 and a minimum read length of 100 bases using Trimmomatic ^66^. Paired-end reads were merged and mapped to a reference sequence using Bowtie2 ^67^. GD/3 and rGD/3 samples were mapped to the consensus sequence of the A/Guangdong/17SF006/2016 human isolate (GISAID isolate ID: EPI_ISL_249309) ^22^. Anhui/1 samples were mapped to the consensus sequence of the A/Anhui/1/2013 human isolate (GISAID isolate ID: EPI_ISL_138739) ^22^. CA04 samples were mapped to A/California/04/2009 reference sequence (GISAID isolate ID: EPI_ISL_29618). To ensure even coverage and reduce resequencing bias, alignment files were randomly subsampled to 200,000 reads per genome using seqtk if total coverage exceeded this value ^68^.

The average genome sequence depth was 40,787 (+/-18,563) reads per genome (**Supplementary Figure 7**). Intrahost single nucleotide variants (iSNVs) were called with Varscan ^69^ using a frequency threshold of 1%, a minimum coverage of 100 reads, and a base quality threshold of Q30 or higher. Variants were called independently for technical replicates and only iSNVs called in both replicates, “intersection iSNVs”, were used for additional analyses ^70^. If an iSNV was only found in one replicate, it was discarded. iSNV frequency is reported as the average frequency found across both replicates. iSNVs are annotated to determine the impact of each variant on the amino acid sequence. iSNVs were annotated in ten open reading frames: PB2 (polymerase basic protein 2), PB1 (polymerase basic protein 1), PA (polymerase acidic), HA (hemagglutinin), NP (nucleoprotein), NA (neuraminidase), M1 (matrix protein 1), M2 (matrix protein 2), NS1 (non-structural protein 1), and NEP (nuclear export protein), though for some analyses M1 and M2 are jointly represented as MP (matrix proteins) an NS1 and NEP are jointly represented as NS (non-structural proteins).

### Sequence data analysis – diversity statistics

Nucleotide diversity was calculated using π summary statistics. π quantifies the average number of pairwise differences per nucleotide site among a set of sequences and was calculated using SNPGenie ^71,72^. SNPGenie adapts the Nei and Gojobori method of estimating nucleotide diversity (π), and its synonymous (π_S_) and nonsynonymous (π_N_) partitions from next-generation sequencing data ^73^. As most random nonsynonymous mutations are likely to be disadvantageous, we expect π_N_ = π_S_ suggests neutrality and that allele frequencies are determined primarily by genetic drift. π_N_ < π_S_ indicates purifying selection is acting to remove new deleterious mutations, and π_N_ > π_S_ indicates diversifying selection is favoring new mutations and may indicate positive selection is acting to preserve multiple amino acid changes ^74^. We used paired t-tests to evaluate the hypothesis that π_N_ = π_S_ within gene segments as well as the hypothesis that π_N_ = π_S_ across samples. Code is available to replicate these analyses in the GitHub repository accompanying this manuscript ^33^.

### Sequence data analysis – estimating transmission bottleneck size

The beta-binomial model, explained in detail in Sobel-Leonard et al ^46^, was used to infer effective transmission bottleneck size (Nb), meaning the number of virions that successfully establish lineages persisting to the first sampling time point in the recipient. In this model, the probability of iSNV transmission is determined by iSNV frequency in the donor at the time of sampling. The probability of transmission is the probability that each iSNV is included at least once in a sample size equal to the bottleneck. The model incorporates sampling noise arising from a finite number of reads and therefore accounts for the possibility of false-negative variants that are not called in recipient animals due to conservative variant-calling thresholds (≥1% in both technical replicates). Code for estimating transmission bottleneck sizes using the beta-binomial approach has been adapted from the original scripts, available here: https://github.com/koellelab/betabinomial_bottleneck.

### Sequence data analysis – enumerating iSNVs occurrences in surveillance samples

H7N9 phylogenies obtained from Nextstrain ^34^ in a JSON format were parsed using a custom python script adapted from Moncla et al ^50^ to extract nonsynonymous amino acid substitutions from the terminal nodes along a phylogenetic tree. We extracted a list of mutations from this tree and associated each mutation with the corresponding host of origin (avian host or human host). We found the intersection between iSNVs detected in our GD/3 and Anhui/1 datasets and the mutations parsed from the phylogenetic tree and counted the number of occurrences each iSNV was found in avian sequences, human sequences, or both. We tested whether occurrences of our iSNVs were enriched in human versus avian datasets using Fisher’s exact test. For readability, the iSNVs represented in **Figure 6** were filtered to remove iSNVs with less than two occurrences in human and/or avian hosts. The complete visualization of the iSNVs and their occurrences are displayed in **Supplementary Figure 5**. Code to replicate these analyses are available in the GitHub repository accompanying this manuscript ^33^.

## Figures

All figures were generated using R (ggplot2) or Python (Matplotlib) with packages including plotly, seaborn, numpy, and scipy and were edited using Adobe Illustrator for clarity and readability. Figure 1 was created using BioRender and Adobe Illustrator. All derived data, raw figure files, and code used to generate the raw figures is available in the GitHub repository accompanying this manuscript ^33^.

## Ethics statement

No animal experiments were specifically performed for this study. We used residual nasal swabs collected from ferrets as part of previously published studies ^17,32^. Animal studies were approved prior to the start of the study by the Institutional Animal Care and Use Committee and performed in accordance with the Animal Care and Use Committee guidelines at the University of Wisconsin-Madison.

## Data availability

Primary data generated and analyzed in this study have been deposited in the Sequence Read Archive under Bioproject ID: PRJNA758865. Individual SRA identifiers can also be found in our GitHub repository.

## Acknowledgements

We would like to acknowledge and thank Dr. Kelsey Florek for her work to develop the Sniffles bioinformatic pipeline for identification of mutations in influenza populations. We also thank Dr. Louise Moncla for her guidance on Figure 6 and help navigating global surveillance data available on Nextstrain. We thank research groups from all over the world for surveilling and sharing their viral sequence data. We also thank the teams at GISAID for maintaining a repository for influenza sequence data and to the Nexstrain team for providing open-source toolkits for analysis and visualization of this data.

## Author Contributions

K.M.B. contributed conceptualization, data curation, formal analysis, investigation, methodology, project administration, software, visualization, writing – original draft preparation, writing – review and editing.

L.A.H. contributed formal analysis, methodology, software, visualization, writing – original draft preparation, writing – review and editing.

C.M.C. contributed project administration, visualization, writing – original draft preparation, writing – review and editing.

G.L.B. contributed investigation, methodology, project administration, writing – review and editing.

J.L. contributed software.

G.N. contributed conceptualization, investigation, writing – review and editing.

T.W. contributed conceptualization, investigation, writing – review and editing.

M.I. contributed conceptualization, investigation, writing – review and editing.

S.Y. contributed conceptualization, investigation, writing – review and editing.

M.I. contributed conceptualization, investigation, writing – review and editing.

Y.K. contributed conceptualization, resources, supervision, writing – review and editing.

T.C.F. contributed conceptualization, funding acquisition, methodology, supervision, writing – review and editing.

## Supplementary figures and tables

1. Supplementary Figure 1: iSNV frequencies in technical replicates
2. Supplementary Figure 2: Count of iSNVs within individual ferrets and over time
3. Supplementary Figure 3: All iSNVs detected in all non-HA gene segments across all virus groups
4. Supplementary Figure 4: Patterns of H1N1 viral genetic diversity within ferret hosts
5. Supplementary Figure 5: iSNVs found in H7N9 global surveillance sequences
6. Supplementary Figure 6: iSNVs in GD/3 inoculum virus
7. Supplementary Figure 7: Sequencing read depth across gene segments
8. Supplementary Table 1: Ferret pairs and transmission time points
9. Supplementary Table 2: iSNV frequency distribution per virus group
10. Supplementary Table 3: Raw data for Figure 6

