## Supplementary_files for "Avian H7N9 influenza viruses are evolutionarily constrained by stochastic processes during replication and transmission in mammals"

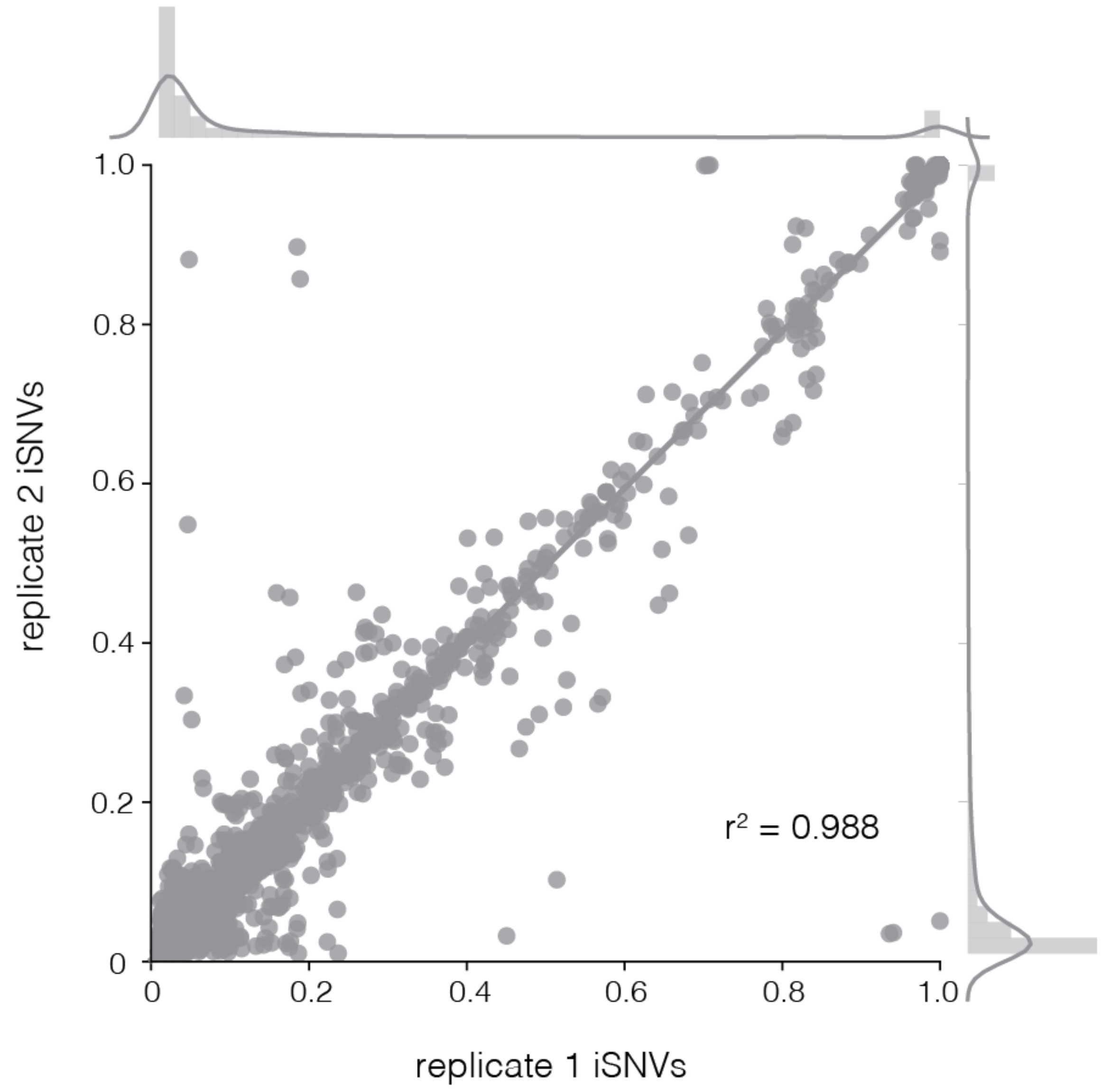


**Supplementary Figure 1: iSNV frequencies in technical replicates**

iSNV frequencies in replicate 1 are shown on the x-axis and replicate 2 are shown on the y-axis. iSNV abundance is plotted along the top and right aspects of the plot using a simple histogram. The r-squared linear correlation is documented in the lower right corner of the plot.

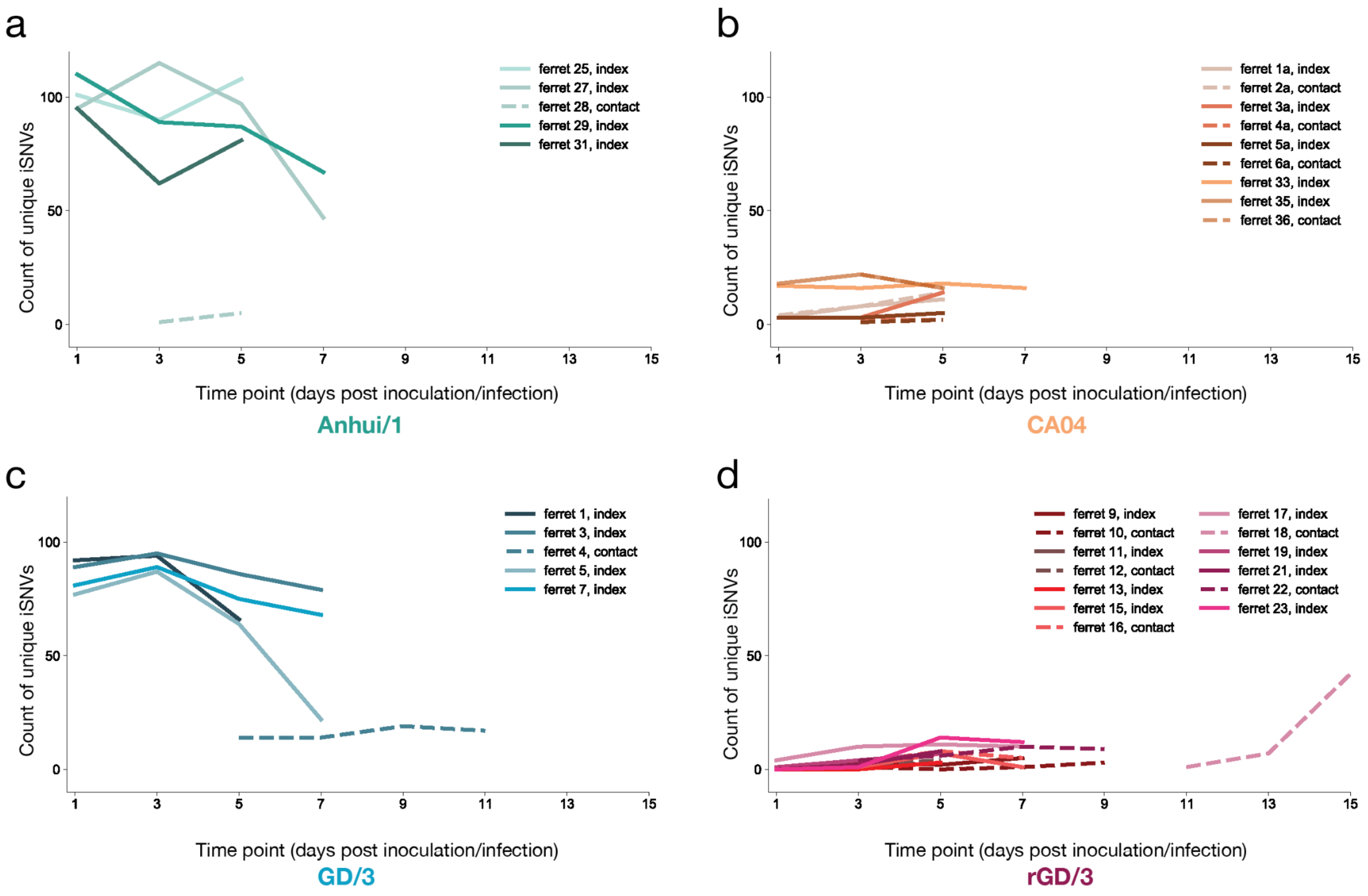


**Supplementary Figure 2: Count of iSNVs within individual ferrets and over time**

iSNVs at different genome sites in **a**. the LPAI H7N9 Anhui/1 group, **b.** the H1N1 CA04 group, **c.** the HPAI H7N9 GD/3 group, and **d.** the recombinant HPAI rGD/3 groups. Donor ferrets are denoted with solid lines and recipient ferrets are denoted with dashed lines.

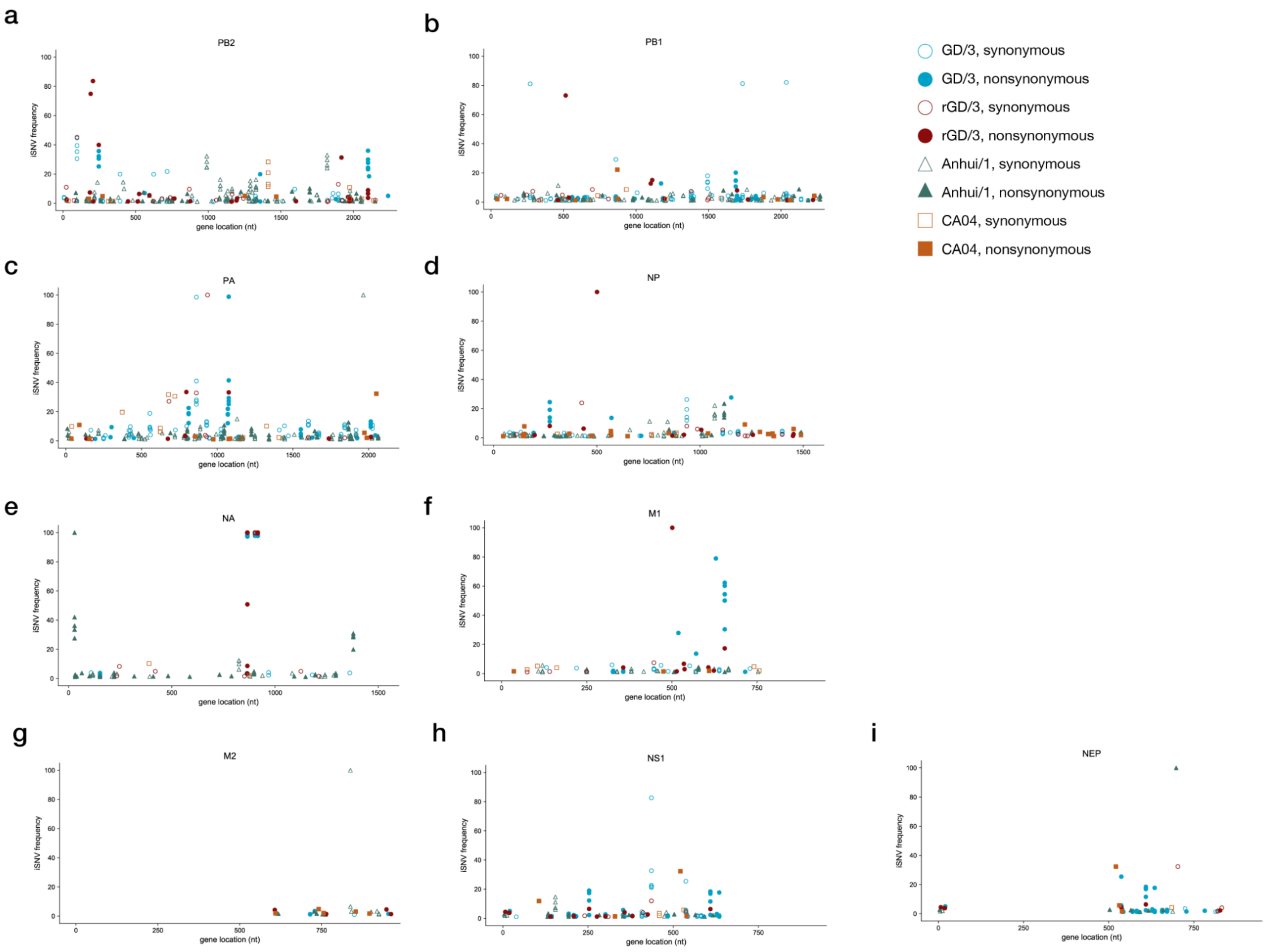


**Supplementary Figure 3: All iSNVs detected in all non-HA gene segments across all virus groups**

All iSNVs detected in **a**. PB2 (polymerase basic protein 2) **b**. PB1(polymerase basic protein 2) **c**. PA (polymerase acidic protein) **d**. NP (nucleoprotein) **e**. NA (neuraminidase) **f**. M1 (matrix protein 1) **g**. M2 (matrix protein 2) **h**. NS1 (non-structural protein 1) **i**. NEP (nuclear export protein or non-structural protein 2). GD/3 and rGD/3 iSNVs are plotted using circles, Anhui/1 iSNVs are plotted using triangles, and CA04 iSNVs are plotted with squares. Synonymous iSNVs are denoted with open symbols and nonsynonymous iSNVs are denoted with closed symbols.

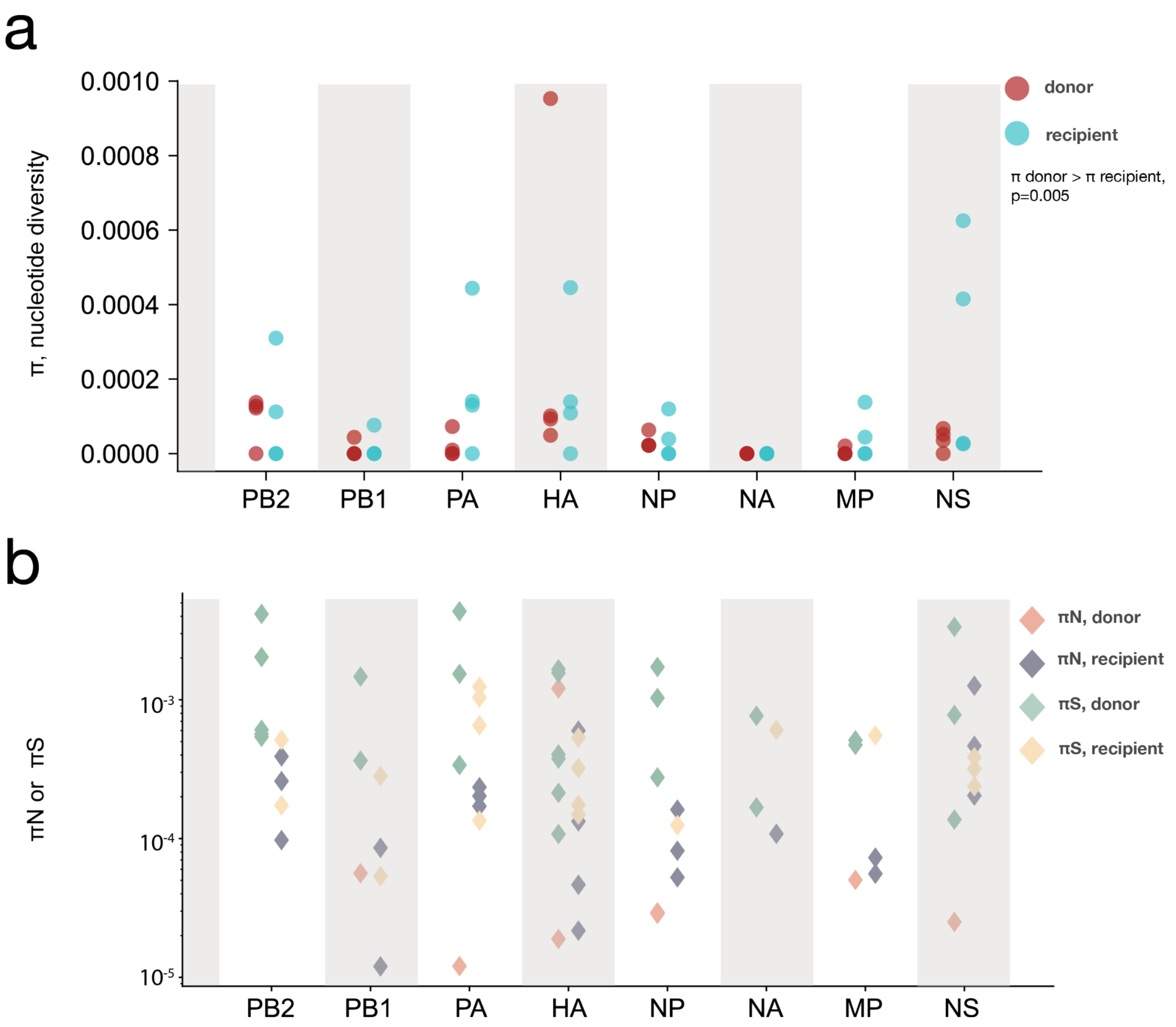


**Supplementary Figure 4: Patterns of H1N1 viral genetic diversity within ferret hosts**

**a**. Genewise nucleotide diversity is plotted for all H1N1 transmission pairs. The donor ferrets are shown in brick-red and the recipient ferrets are shown in aqua-blue. Nucleotide diversity does not significantly differ between donor and recipient ferrets in any single gene segment or at the genome level ($p=0.278$; paired t-test). **b**. πN and πS in the H1N1 donors and recipients are plotted for each gene segment. πN and πS in the donor ferrets are denoted by the salmon and green diamonds, respectively. πN and πS in the recipient ferrets are denoted by the dark blue and yellow diamonds, respectively.

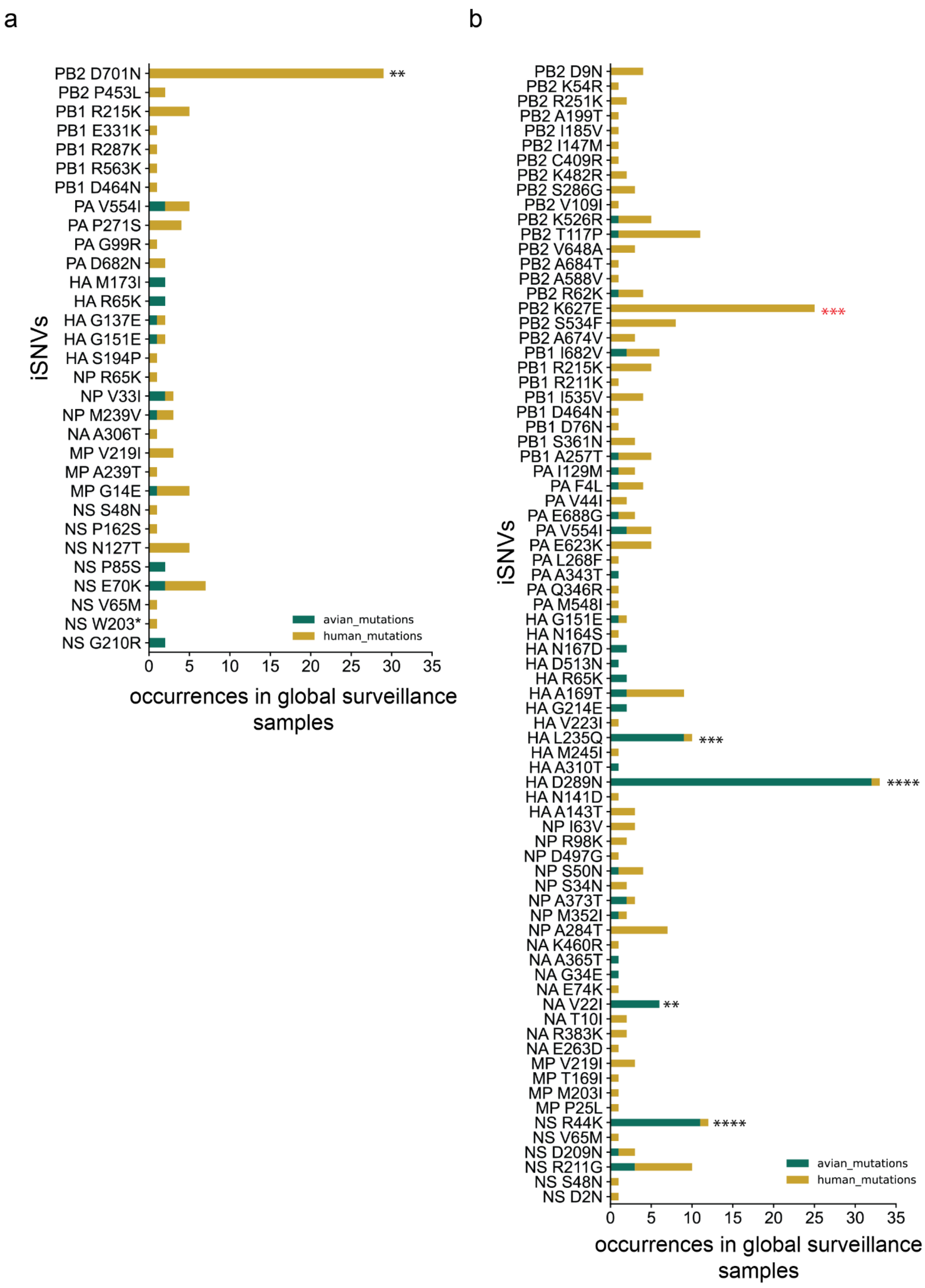


**Supplementary Figure 5: iSNVs found in H7N9 global surveillance sequences**

Occurrences of iSNVs in H7N9 global surveillance sequences in the (**a**) GD/3, (**b**) and Anhui/1 datasets. Fisher's exact test was used to assess for significant enrichment in human or bird sequences and these results are shown with asterisks (* p = 0.05 - 0.01, ** p = 0.01 - 0.001, *** p = 0.0001 - 0.0000).

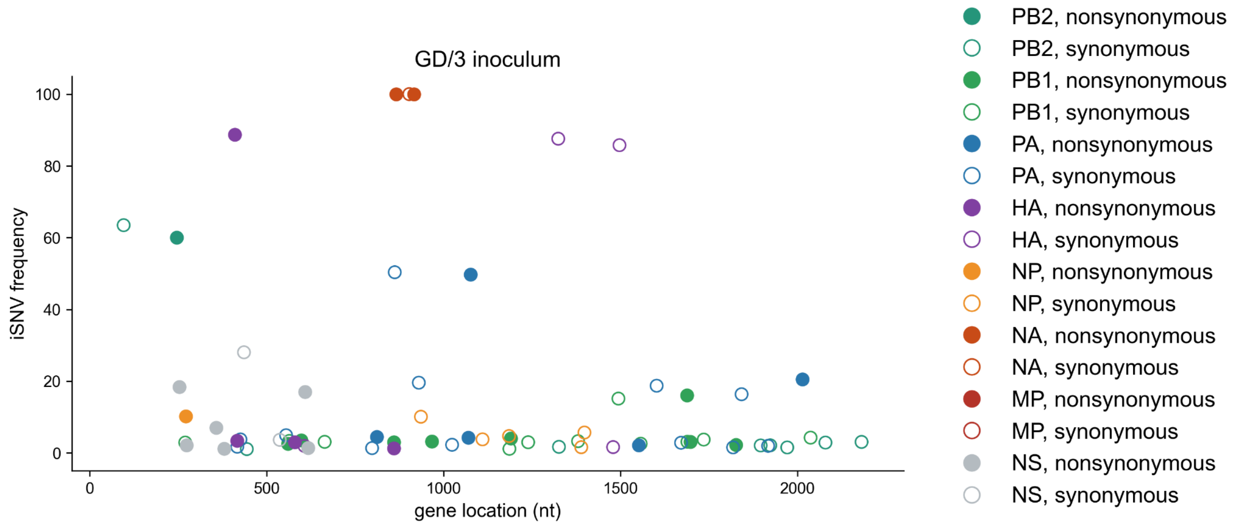


**Supplementary Figure 6: iSNVs in GD/3 inoculum virus**

iSNVs in the GD/3 inoculating virus are plotted with frequency (%) on the y-axis and gene location (nucleotide numbering) on the x-axis. Synonymous iSNVs are denoted with open symbols and nonsynonymous iSNVs are denoted with closed symbols. iSNVs found in PB2 are shown in turquoise, PB1 in green, PA in blue, HA in purple, NP in light orange, NA in dark orange, MP in red, and NS in gray. Note that each gene's numbering starts at x=0 and the gene segments are not plotted sequentially.

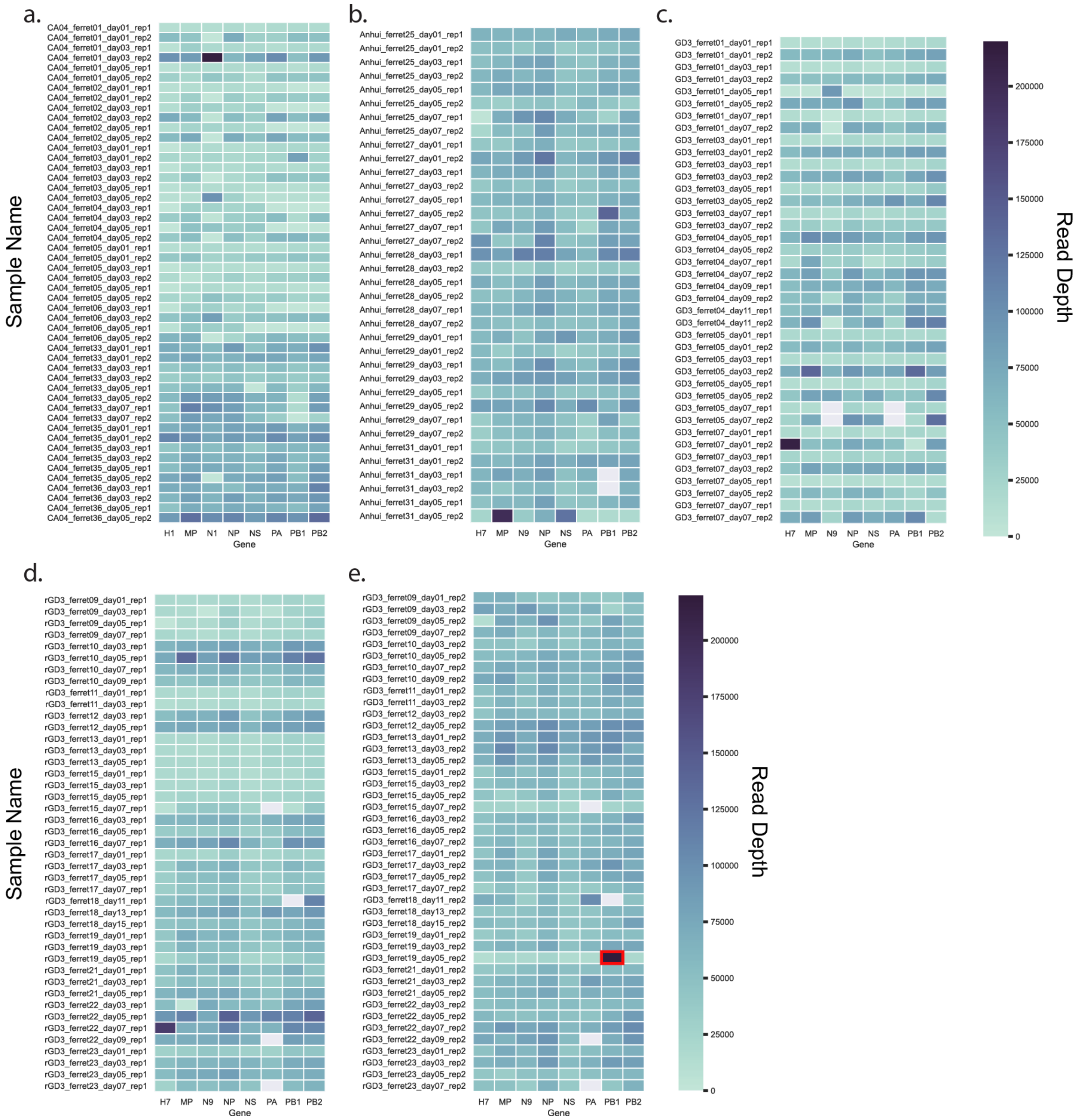


**Supplementary Figure 7: Sequencing read depth across gene segments**

Each row represents an individual nasal wash sample collected from a ferret at specific timepoints for both technical replicates. Each of the 8 columns represent each gene segment: H1/H7 (hemagglutinin subtypes 1 and 7), MP (matrix protein), N1/N9 (neuraminidase subtypes 1 and 7), NP (nucleoprotein), NS (non-structural protein), PA (polymerase acidic), and PB1/PB2 (polymerase basic proteins 1 and 2). Rectangles shows sequencing read depth for **a.** A/California/04/2009 (CA04), **b.** A/Anhui/1/2013 (Anhui), **c.** A/Guangdong/17SF003/2016 (GD3) **d.** A/Guangdong/17SF003/2016 rGD3-NA289R and **e.** A/Guangdong/17SF003/2016 rGD3-NA289K. Highlighted depth (red rectangle) shows an outlier with read depth to 748,314, while the remaining samples ranged from a value of 0 up to 216,190 (in addition to missing values shown in gray rectangles).

**Supplementary Table 1: Ferret pairs and transmission time points**

Ferret identification numbers for transmitting and non-transmitting pairs. First available time point of the recipient ferret following transmission can be found on the last column.

| | | Ferret ID | | First available timepoint for recipient |
|---|---|---|---|---|
| | | Infected (Donor) | Contact (Recipient) | |
| **CA04** | | **1** | **2** | **Day 1** |
| | | **3** | **4** | **Day 3** |
| | | **5** | **6** | **Day 3** |
| | | **33** | **36** | **Day 3** |
| | | 35 | - | - |
| **Anhui/1** | | 25 | - | - |
| | | **27** | **28** | **Day 3** |
| | | 29 | - | - |
| | | 31 | - | - |
| Guangdong | **GD/3** | 1 | - | - |
| | | **3** | **4** | **Day 5** |
| | | 5 | - | - |
| | | 7 | - | - |
| | **rDG/3 NA-289R** | **9** | **10** | **Day 3** |
| | | **11** | **12** | **Day 3** |
| | | 13 | - | - |
| | | **15** | **16** | **Day 3** |
| | **rGD/3 NA-289K** | **17** | **18** | **Day 11** |
| | | 19 | - | - |
| | | **21** | **22** | **Day 3** |
| | | 23 | - | - |

| **Transmission pair** | Not Transmitted |
|---|---|

**Supplementary Table 2: iSNV frequency distribution per virus group**

The average number of iSNVs per frequency bin within each virus group. Averages and standard deviations are calculated across ferrets within each virus group.

| | Neutral expectation | Anhui/1 | | CA04 | | GD/3 | | rGD/3 | |
|---|---|---|---|---|---|---|---|---|---|
| **bins** | | average | st-dev | average | st-dev | average | st-dev | average | st-dev |
| **1-10%** | 0.501094 | 0.859606 | 0.075788 | 0.636087 | 0.167351 | 0.624697 | 0.139581 | 0.816804 | 0.258179 |
| **10-20%** | 0.150844 | 0.081222 | 0.044959 | 0.178585 | 0.126434 | 0.112123 | 0.026236 | 0.032326 | 0.046395 |
| **20-30%** | 0.088238 | 0.030318 | 0.016055 | 0.128289 | 0.151903 | 0.054854 | 0.027102 | 0.062271 | 0.127024 |
| **30-40%** | 0.062606 | 0.018988 | 0.010798 | 0.013538 | 0.023612 | 0.047376 | 0.028516 | 0.01946 | 0.035594 |
| **40-50%** | 0.048561 | 0.006027 | 0.005327 | 0.018849 | 0.033898 | 0.020635 | 0.005861 | 0.003053 | 0.007326 |
| **50-60%** | 0.039677 | 0.001077 | 0.001321 | 0.016339 | 0.030944 | 0.030021 | 0.038061 | 0.014957 | 0.03198 |
| **60-70%** | 0.033547 | 0.000617 | 0.001235 | 0.008312 | 0.012043 | 0.013663 | 0.013572 | 0.009615 | 0.033309 |
| **70-80%** | 0.029059 | 0.000823 | 0.001646 | 0 | 0 | 0.023043 | 0.028008 | 0.019231 | 0.066617 |
| **80-90%** | 0.025632 | 0 | 0 | 0 | 0 | 0.018948 | 0.012351 | 0.021062 | 0.06639 |
| **90-99%** | 0.020742 | 0.001321 | 0.002642 | 0 | 0 | 0.05464 | 0.068023 | 0.001221 | 0.00423 |

**Supplementary Table 3: Raw data for Figure 6**

Summary of the number of total iSNVs and iSNVs-per-gene that we identified in our Anhui/1 and GD/3 datasets that were also identified in Nextstrain surveillance samples.

| | | Anhui/1 | | | | | GD/3 | | | | |
|---|---|---|---|---|---|---|---|---|---|---|---|
| **Gene** | **Num. Nextstrain mutations** | **Num. of iSNVs** | **Num. of iSNVs found on Nextstrain sequences** | **Num. of iSNVs found on Human sequences** | **Num. of iSNVs found on Avian sequences** | **Num. of iSNVs found on Human and Avian sequences** | **Num. of iSNVs** | **Num. of iSNVs found on Nextstrain sequences** | **Num. of iSNVs found on Human sequences** | **Num. of iSNVs found on Avian sequences** | **Num. of iSNVs found on Human and Avian sequences** |
| **HA** | 405 | 12 | 5 | 1 | 2 | 2 | 24 | 14 | 5 | 5 | 4 |
| **NA** | 216 | 3 | 1 | 1 | 0 | 0 | 19 | 8 | 5 | 3 | 0 |
| **PA** | 303 | 14 | 5 | 3 | 1 | 1 | 23 | 10 | 5 | 1 | 4 |
| **NP** | 220 | 23 | 4 | 1 | 1 | 2 | 19 | 8 | 5 | 0 | 3 |
| **MP** | 106 | 6 | 4 | 2 | 1 | 1 | 11 | 4 | 4 | 0 | 0 |
| **NS** | 207 | 15 | 8 | 5 | 2 | 1 | 14 | 6 | 3 | 0 | 3 |
| **PB1** | 314 | 15 | 5 | 5 | 0 | 0 | 20 | 8 | 6 | 0 | 2 |
| **PB2** | 312 | 9 | 3 | 2 | 1 | 0 | 35 | 19 | 16 | 0 | 3 |
| **Total SNV's** | 2083 | 97 | 35 | 20 | 8 | 7 | 165 | 77 | 49 | 9 | 20 |
